## Supplemental Materials for "Data-driven, participatory characterization of farmer varieties discloses teff breeding potential under current and future climates"

### Figure Supplements

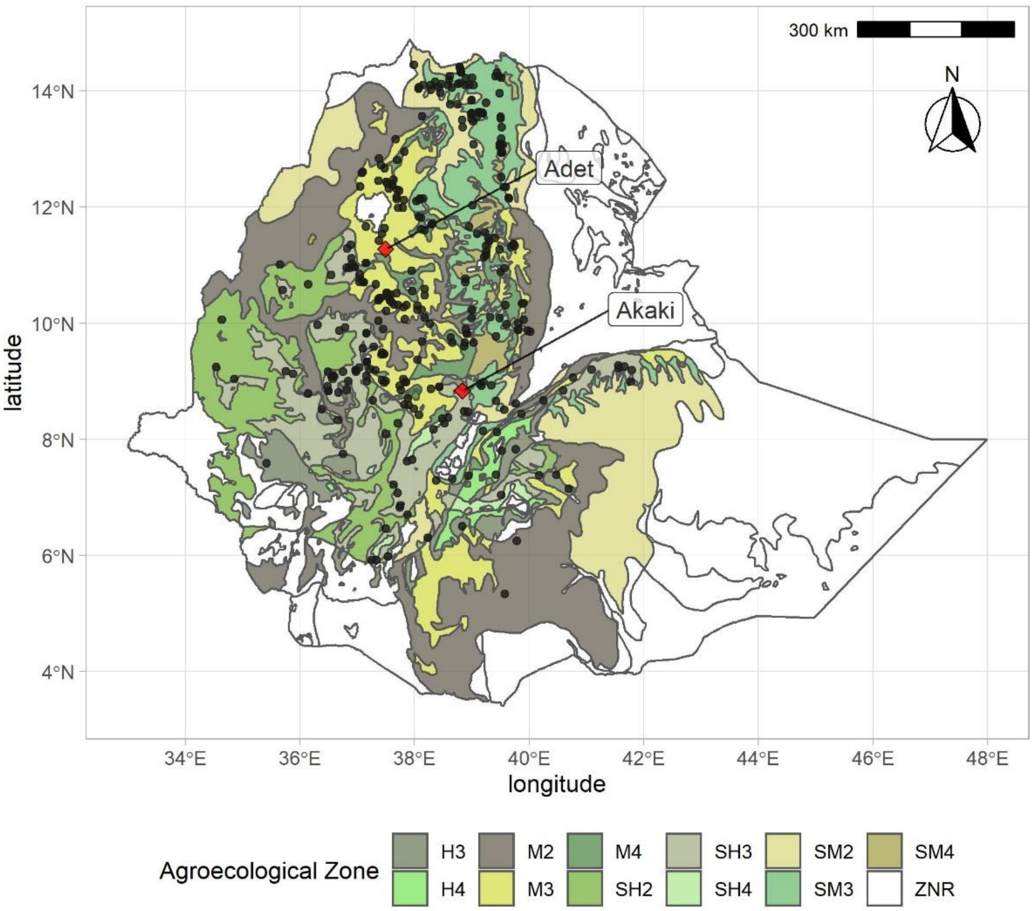


**Figure 1–figure supplement 1**. Distribution of georeferenced farmer varietis in the EtDP (N= 314) overlaid to agroecological zones of Ethiopia. Sampling locations are indicated with black points. Adet and Akaki, the two field experiment sites, are indicated with red diamonds. H3, tepid humid mid-highlands; H4, cool humid mid-highlands; M2, warm moist lowlands; M3, tepid moist mid-highlands; M4, cool moist mid-highlands; SH2, warm sub-humid lowlands; SH3, tepid sub-humid mid-highlands; SH4, cool sub-humid mid-highlands; SM2, warm sub-moist lowlands; SM3, tepid sub-moist mid-highlands; SM4, cool sub-moist mid-highlands; ZNR, zone not relevant (no hits in the teff collection).

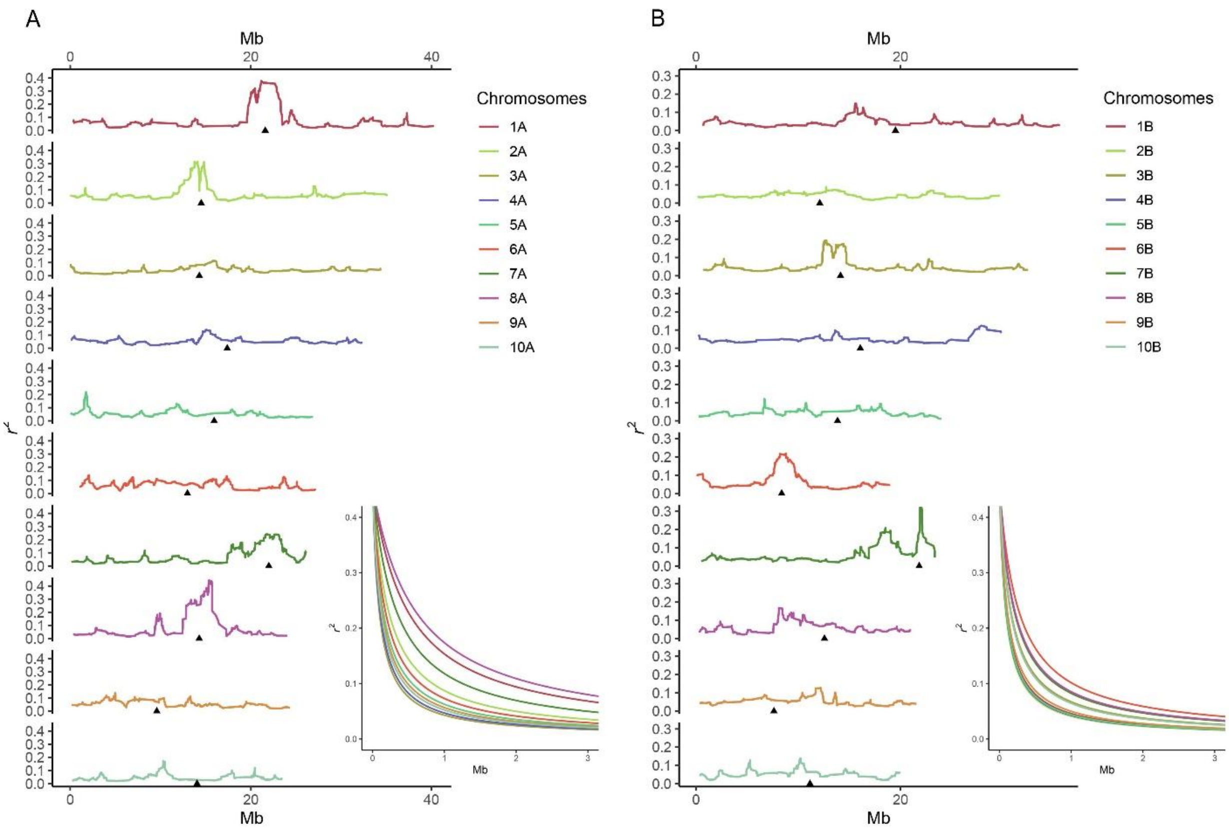


**Figure 1–figure supplement 2**. Linkage disequilibrium (LD) in subgenome A (**A**) and subgenome B (**B**) calculated as $r^2$. Each panel shows LD evolution in the ten chromosomes of each subgenome, from number 1 to the top to number 10 to the bottom. The lines represent LD measures as a rolling window across the chromosome and are colored according to legend. Black triangles represent centromeres. The inset represents LD decay as a function of physical distance of markers. Mb, million basepairs.

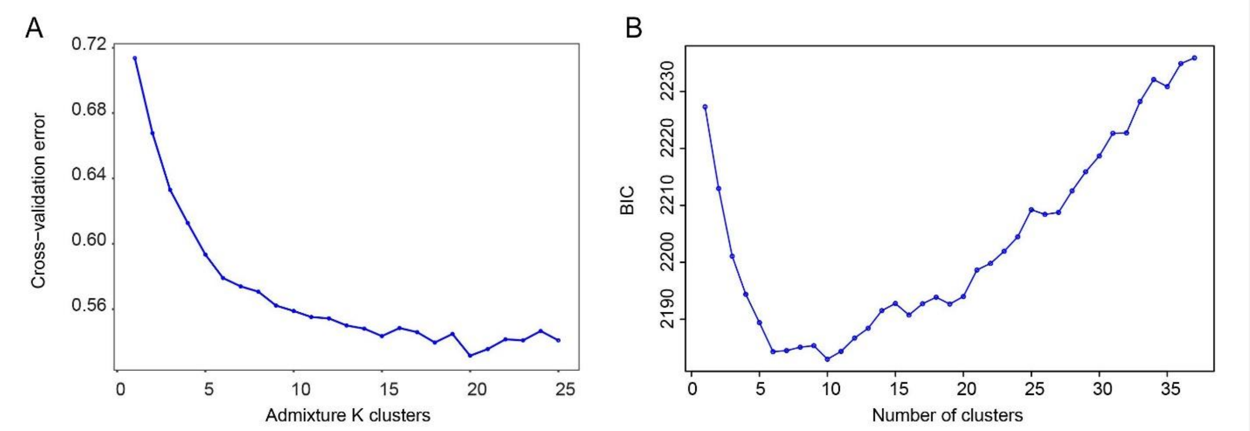


**Figure 1–figure supplement 3**. Predictive accuracy of the model-based unsupervised clustering (ADMIXTURE) and the Discriminant Analysis of Principal Components (DAPC) using the 5-fold cross-validation procedure and the Bayesian Information Criterion (BIC), respectively. (**A**) Cross-validation (CV) error for Admixture clusters ranging from K=2 to K=25. (**B**) BIC values for different values of DAPC clusters. The optimal number of clusters is 10.

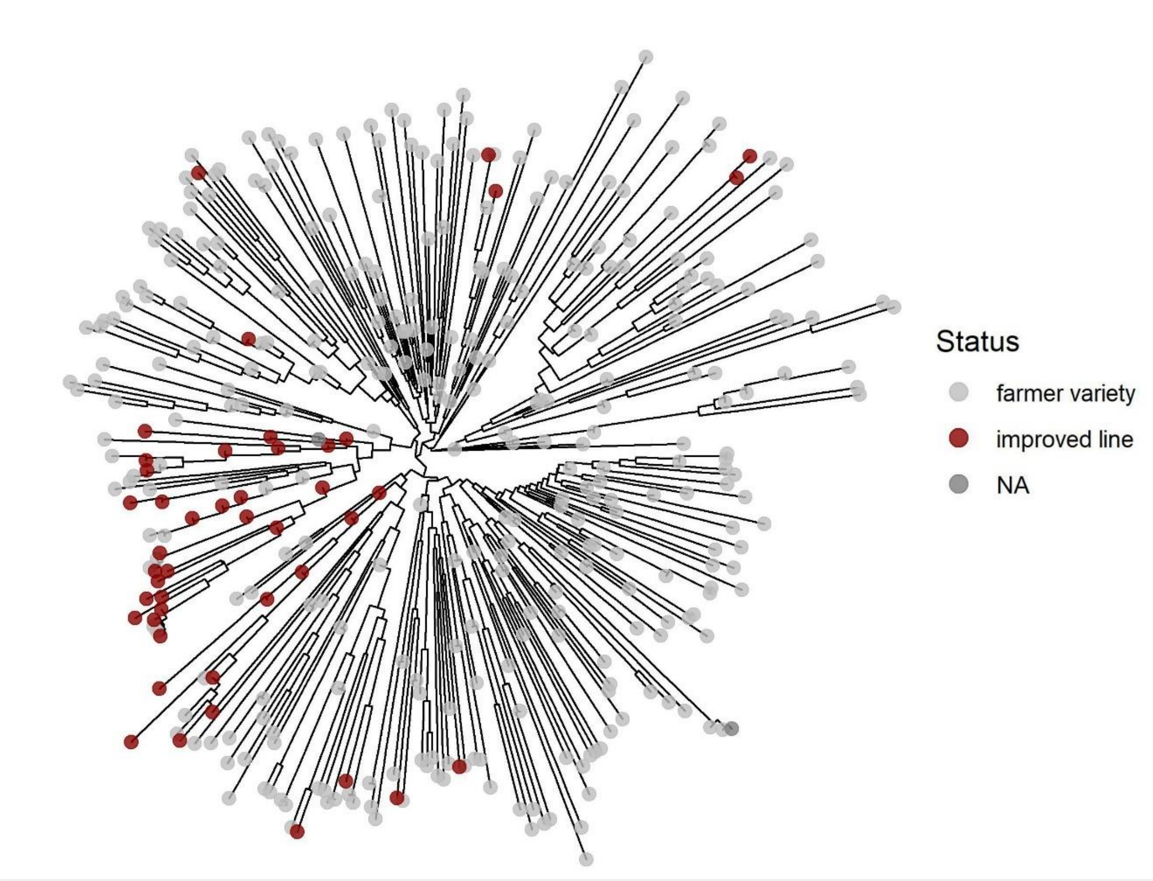


**Figure 1–figure supplement 4.** Unrooted neighbor-joining phylogenetic trees of the EtDP (N=366). Taxa are color coded according their status as in the legend.

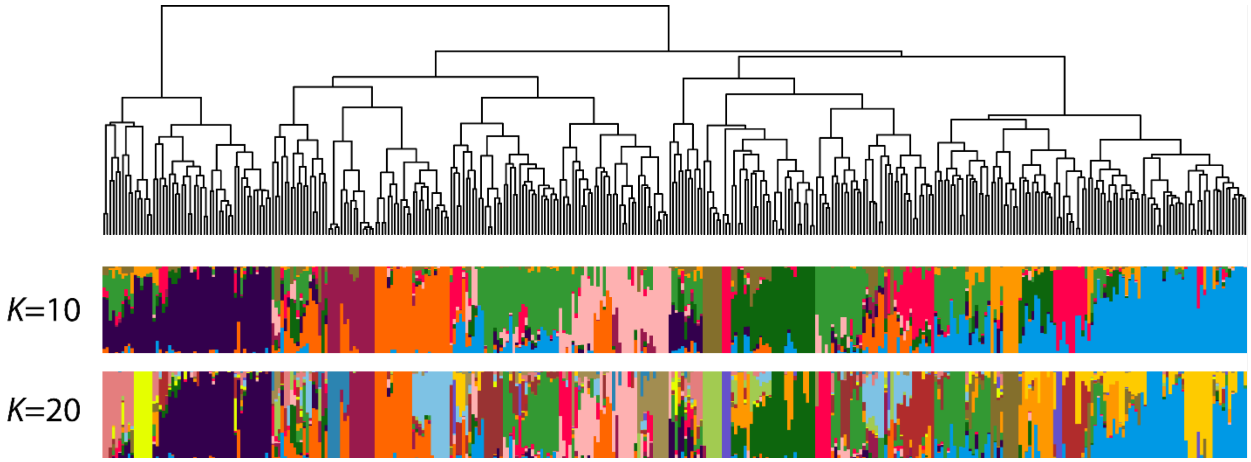


**Figure 1–figure supplement 5**. Population structure of the core collection (N=366). The dendrogram represents the result of IBS-based hierarchical clustering analysis. Each individual is represented by a vertical bar, colored according to its belonging to one of the predicted clusters. The two bar plots represent ADMIXTURE assignation for a value of K=10 (corresponding to the best DAPC interpretation) and K=20. Accessions on the *x*-axis are ordered according to the dendrogram on top.

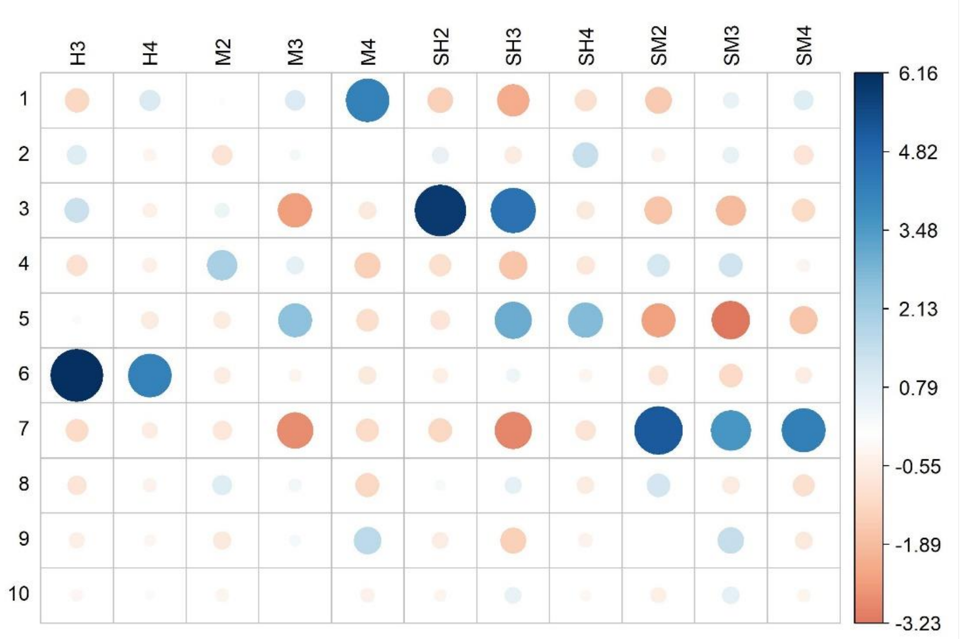


**Figure 1–figure supplement 6**. Residual plot for the Pearson's Chi-squared test of independence between DAPC genetic clusters and Ethiopian agroecological zones (df=60, p-value < 2.2e-16). Circle colors and sizes indicate the relative contribution of each cell to the Chi-square score. Dark blue and larger circles indicate higher effect sizes. H3, tepid humid mid-highlands; H4, cool humid mid-highlands; M2, warm moist lowlands; M3, tepid moist mid-highlands; M4, cool moist mid-highlands; SH2, warm sub-humid lowlands; SH3, tepid sub-humid mid-highlands; SH4, cool sub-humid mid-highlands; SM2, warm sub-moist lowlands; SM3, tepid sub-moist mid-highlands; SM4, cool sub-moist mid-highlands; ZNR, zone not relevant (no hits in the teff collection).

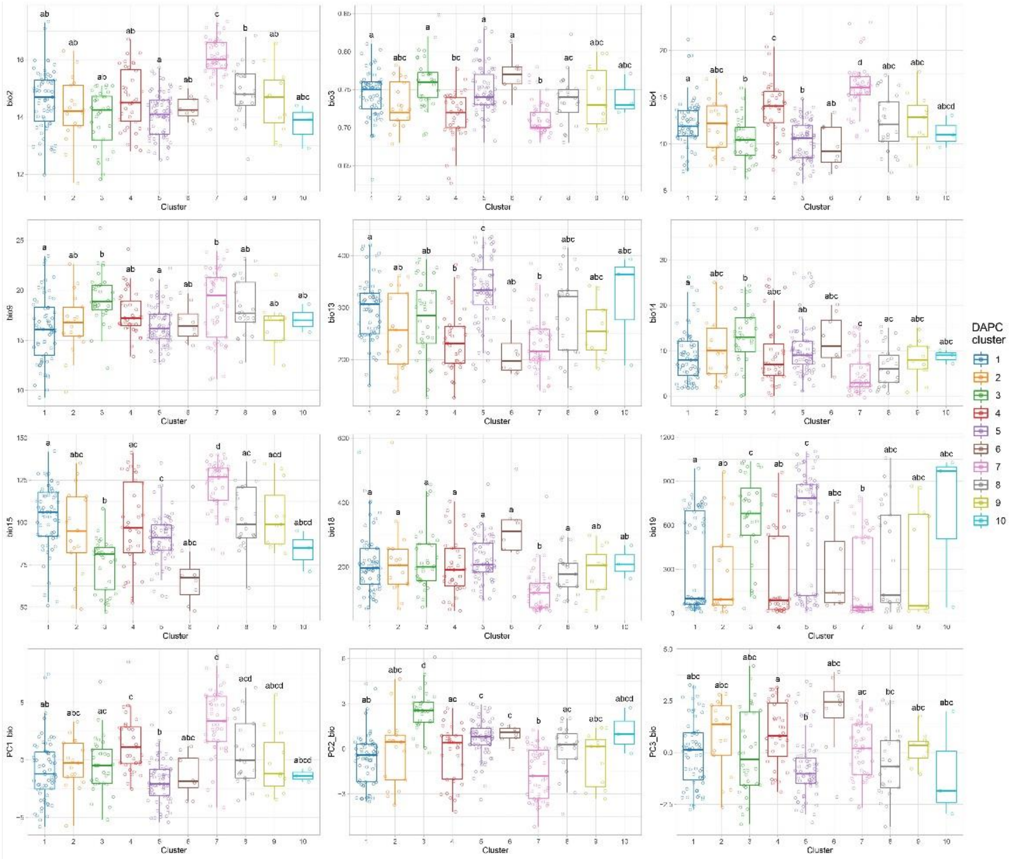


**Figure 2–figure supplement 1**. Bioclimatic differences among the 10 DAPC clusters. Comparisons among clusters were performed using the pairwise Wilcoxon rank sum test with Bonferroni correction for multiple testing. Letters on top of boxplots denote significance levels, with same letters indicating non-significant differences. Only non-collinear bioclimatic variables are shown. Bio2: mean diurnal temperature range; Bio3: Isothermality; Bio4: Temp. Seasonality; Bio9: Mean Temp. Of Driest Quarter; Bio13: Precipitation of Wettest Month; Bio14: Precipitation of Driest Month; Bio15: Precipitation Seasonality; Bio18: Precipitation of Warmest Quarter; Bio19: Precipitation of Coldest Quarter. PC1_bio: first bioclimatic principal component; PC2_bio: second bioclimatic principal component; PC3_bio: third bioclimatic principal component.

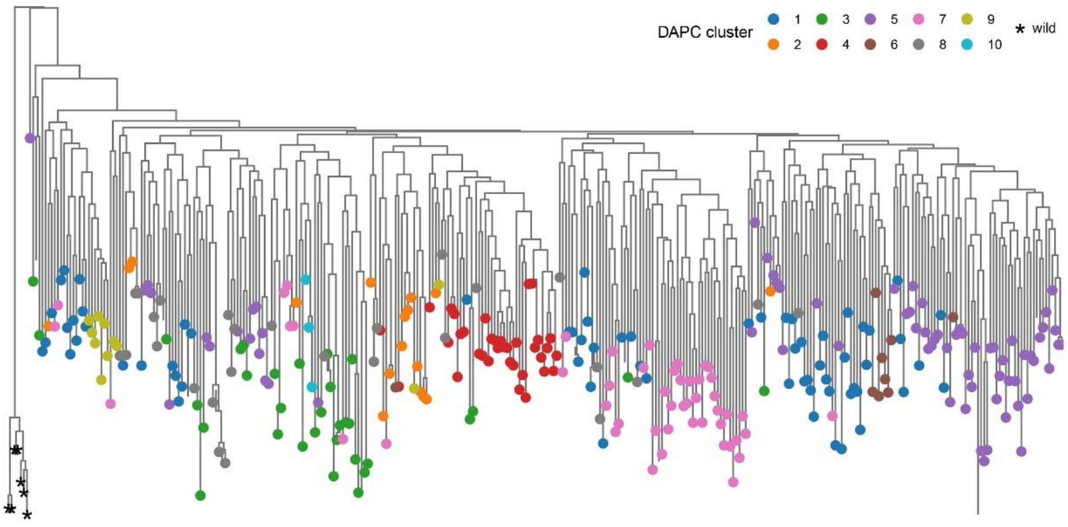


**Figure 2–figure supplement 2**. Neighbor-joining phylogenetic trees of the EtDP rooted with wild relative accessions *Eragrostis pilosa* and *Eragrostis curvula*. Tree tips are colored by either DAPC genetic clusters. Wild relatives are reported to the left of the tree, marked with star signs.

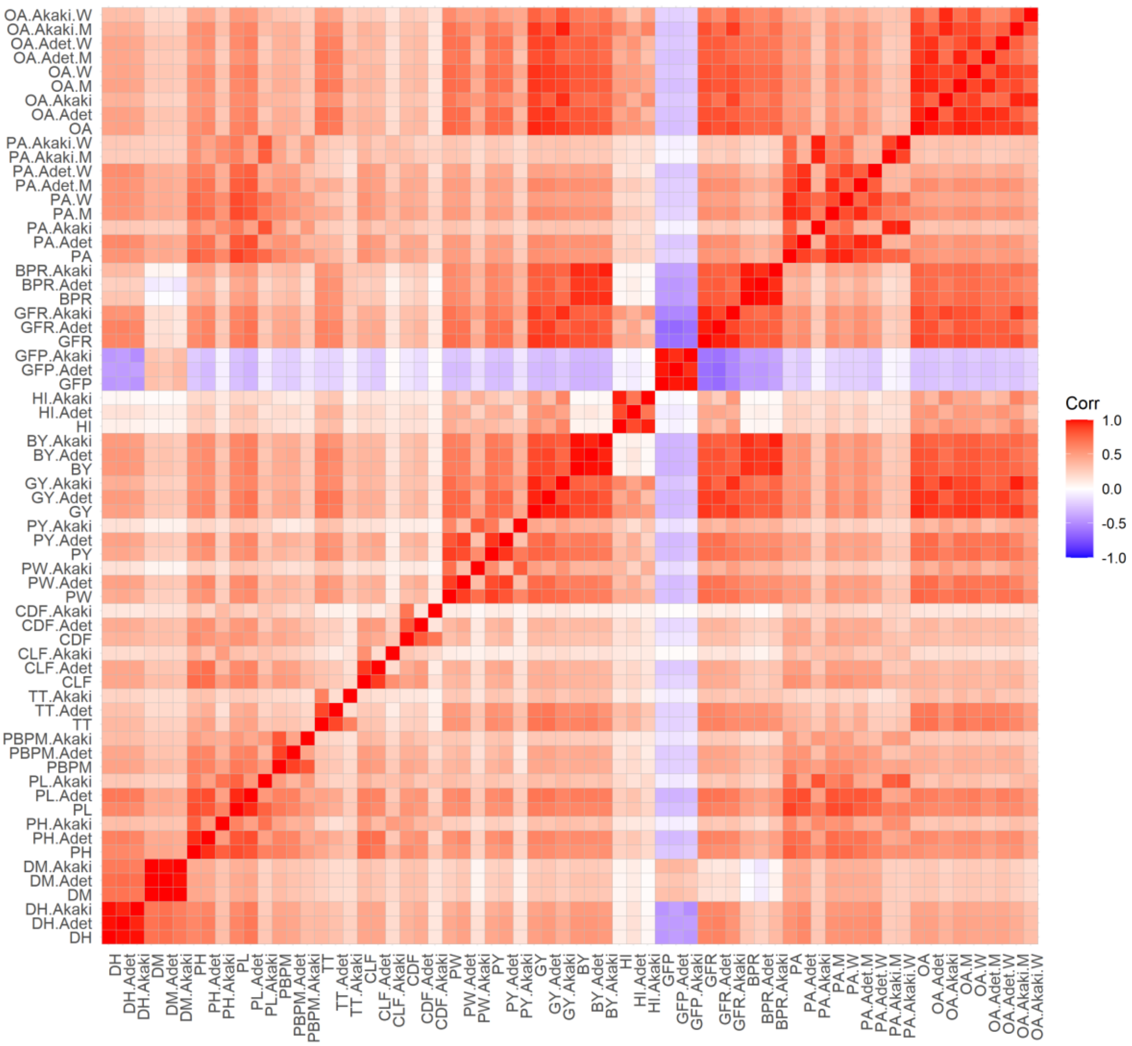


**Figure 3–figure supplement 1**. Correlations between agronomic traits and PVS traits, by location and gender. PVS traits are grouped to the top and right of the plot. The pairwise matrix in the plot represents coefficients of correlation according to the legend to the right. When specific by location, the trait code is attached to either Akaki or Adet. When specific by gender, M (men) or W (women) is attached. DH, days to heading; DM, days to maturity; PH, plant height; PL, panicle length; PBPM, number of primary branches per main shoot panicle; TT, total tillers; CLF, first culm length; CDF, first culm diameter; PW, panicle weight; PY, panicle yield; GY, grain yield; BY, biomass yield; HI, harvest index; GFP, grain filling period; GFR, grain filling rate; BPR, biomass production rate; OA, overall appreciation; PA, panicle appreciation.

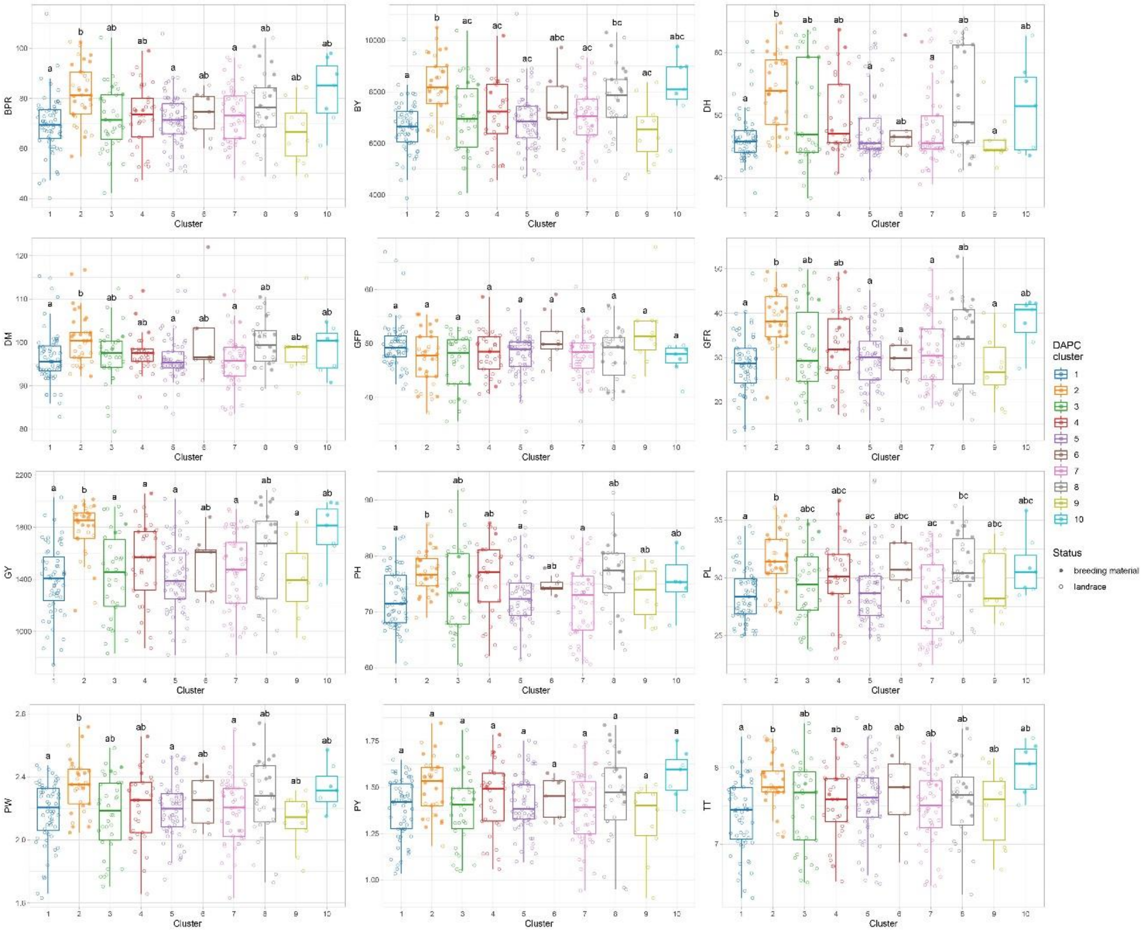


**Figure 3–figure supplement 2**. Phenotypic differences among the ten DAPC clusters. Comparisons among clusters were performed using the pairwise Wilcoxon rank sum test with Bonferroni correction for multiple testing. Letters on top of boxplots denote significance levels, with same letters indicating non-significant differences. Only traits with significant differences are shown. DH, days to heading; DM, days to maturity; PH, plant height; PL, panicle length; PBPM, number of primary branches per main shoot panicle; TT, total tillers; CLF, first culm length; CDF, first culm diameter; PW, panicle weight; PY, panicle yield; GY, grain yield; BY, biomass yield; HI, harvest index; GFP, grain filling period; GFR, grain filling rate; BPR, biomass production rate; OA, overall appreciation; PA, panicle appreciation.

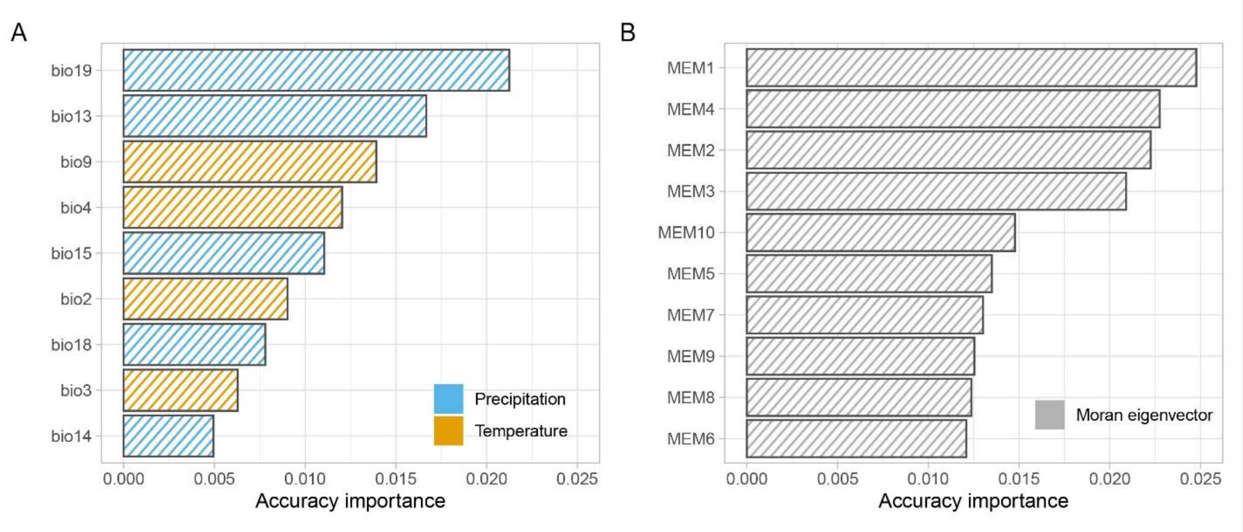


**Figure 4–figure supplement 1**. Ranked accuracy and importance of bioclimatic (**A**) and geographic (MEM) (**B**) variables in predicting turnover in allele frequency using gradient forest analysis. Genomic composition is best predicted by MEMs one to four and precipitation variables (bio19 and bio13).

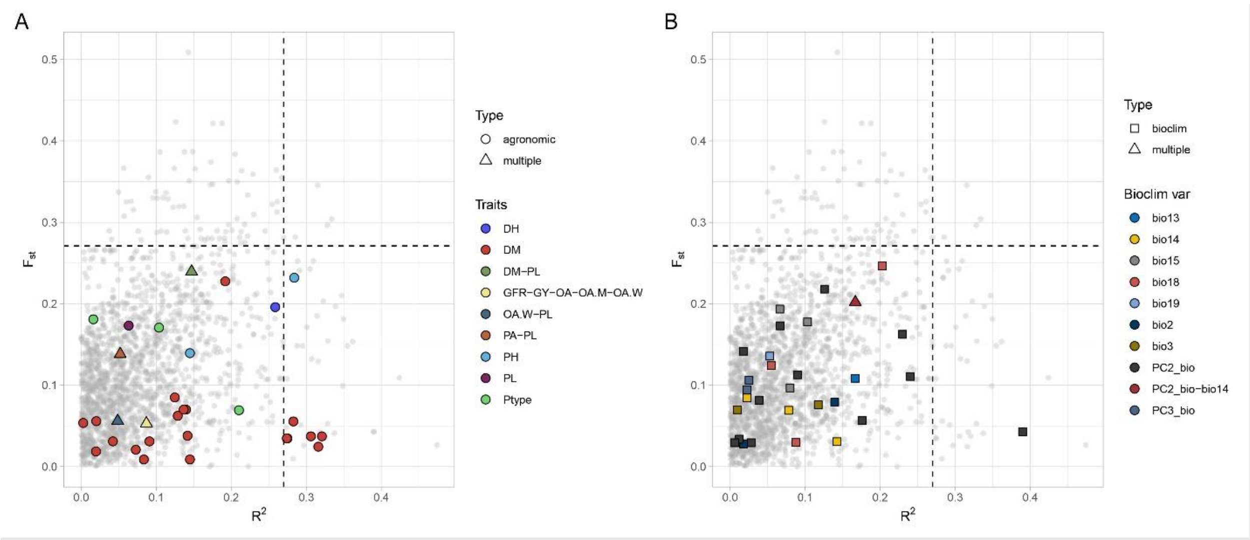


**Figure 4–figure supplement 2.** LD blocks distribution in relation to $F_{st}$ (y axis) and GF $R^2$ values (x axis). LD blocks with null $R^2$ values towards the GF are not shown. LD blocks containing quantitative trait nucleotides (QTNs) are highlighted with colors according to legend. Panel (**A**) reports agronomic and farmer traits, panel (**B**) reports QTNs with bioclimatic traits. In both panels, shapes denote whether QTN target a single trait or rather multiple traits. DH, days to heading; DM, days to maturity; PH, plant height; PL, panicle length; GY, grain yield; GFR, grain filling rate; Ptype, panicle type; PA, panicle appreciation; OA, overall appreciation; OA.W, overall appreciation women; OA.M, overall appreciation men.

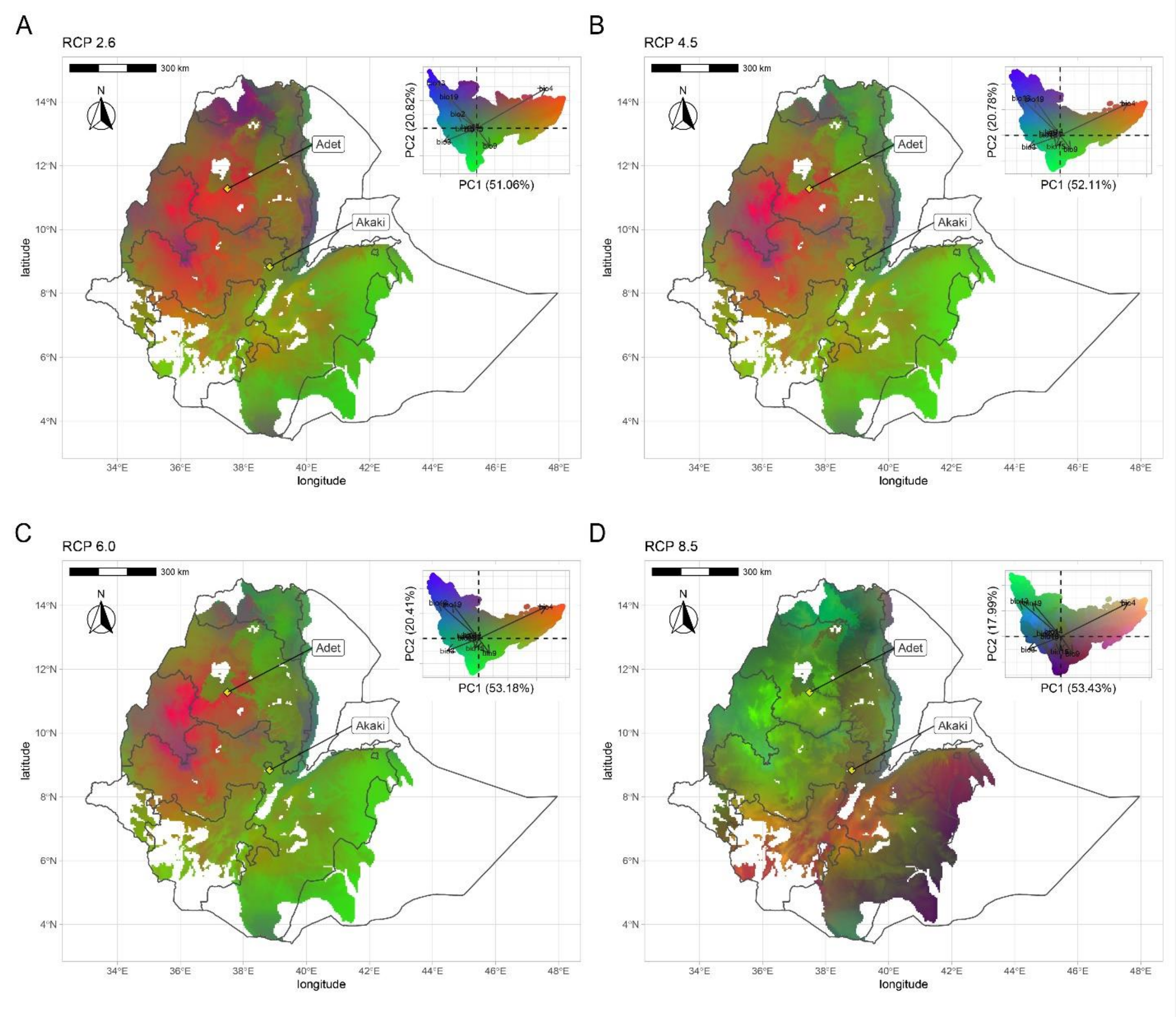


**Figure 4–figure supplement 3.** Geographic distribution of climate-driven allelic variation under four Representative Concentration Pathways: RCP2.6, RCP4.5, RCP 6.0 and RCP8.5. Colors are based on principal components analysis (PCA) of transformed climate variables where red is defined by values of PC1 + PC2, green by negative values of PC2, and blue by PC3+PC2-PC1 (as in the reference manual). Similar colors in the maps represent similar allele frequencies at climate-responsive loci.

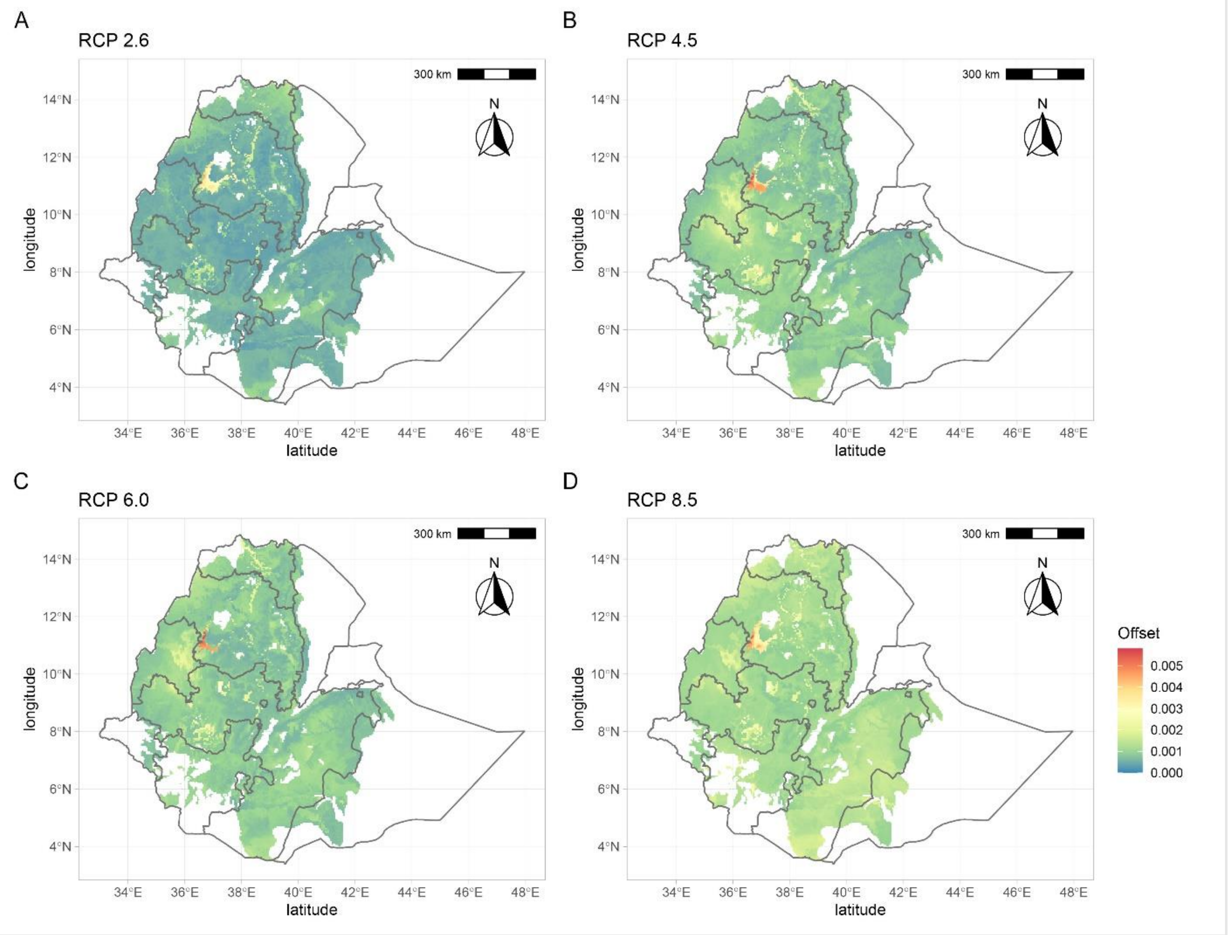


**Figure 4–figure supplement 4.** Genomic vulnerabilities in the teff cropping area based on projections for four representative concentration pathways: RCP2.6, RCP4.5, RCP 6.0 and RCP8.5. The color scale indicates the magnitude of the mismatch (*i.e.* Euclidean distance) between current and projected climate-driven turnover in allele frequencies.

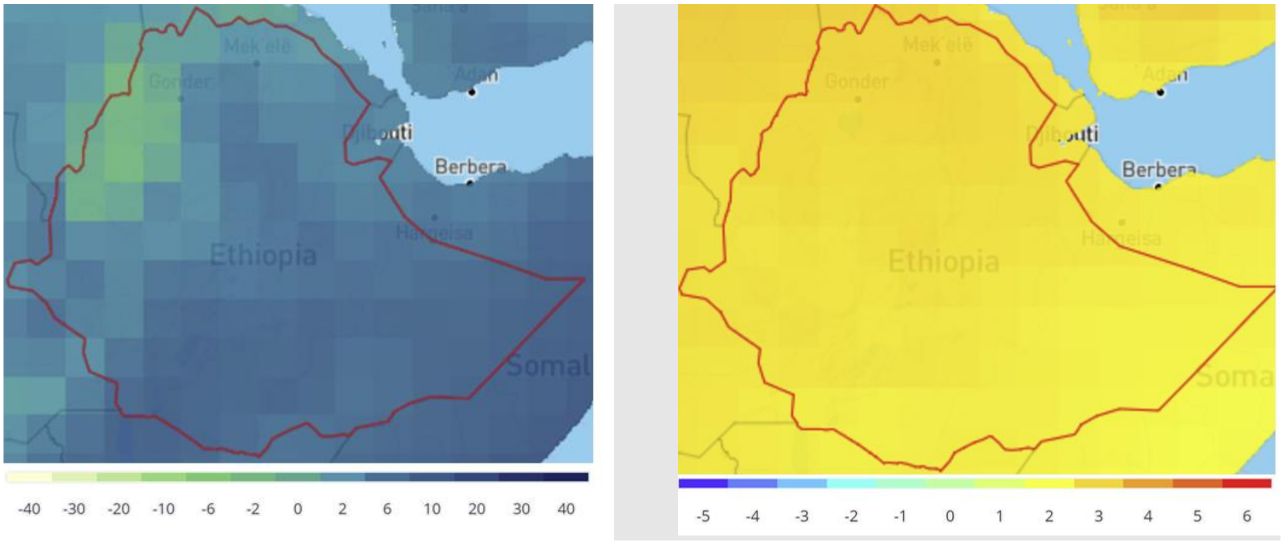


**Figure 4–figure supplement 5.** Projected change in Ethiopian climate for 2070s under RCP8.5 compared to 1986-2005. The plots show the ensemble monthly mean change over 12 months in precipitation (mm) (left) and temperature (°C) (right).

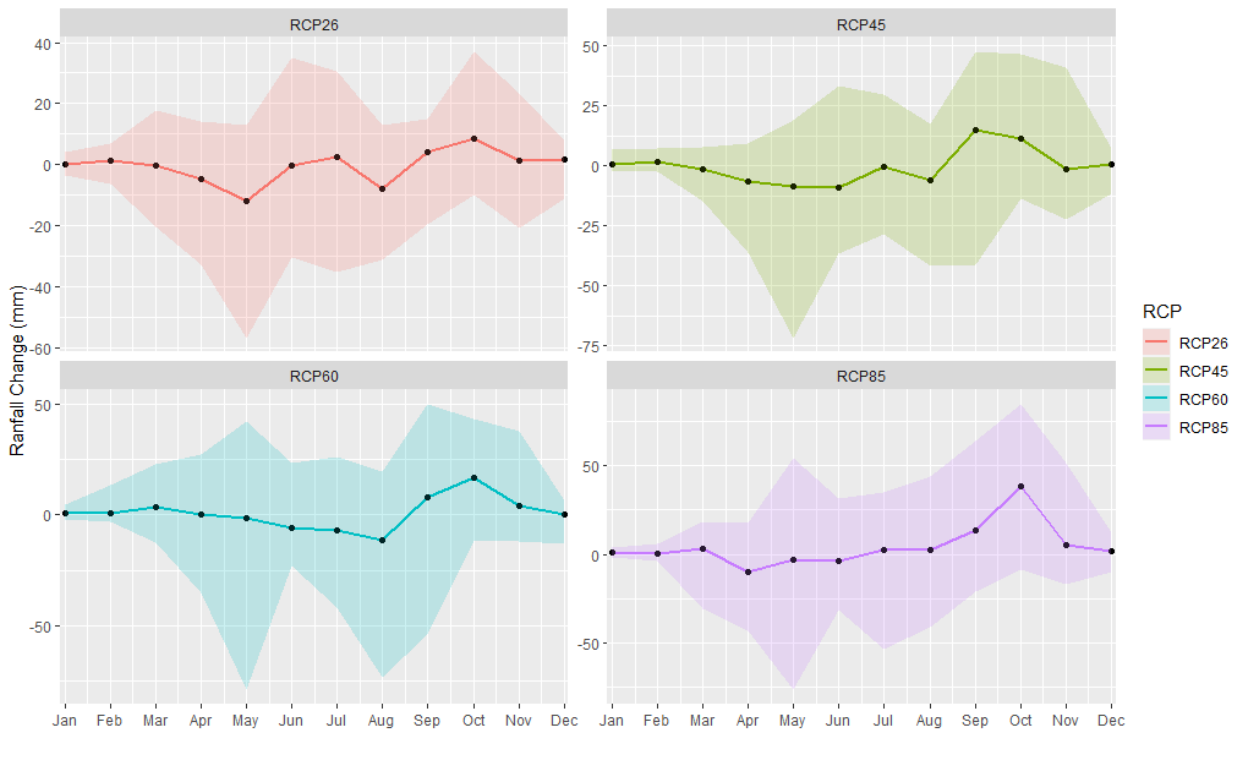


**Figure 4–figure supplement 6.** Mean change in monthly rainfall compared to the reference period in the south lake tana (36.5-37.75 east,10.7- 12 north) by 2070s under RCP2.6 (low emissions), RCP4.5 (medium low-emission), RCP6.0 (medium-high emission), and RCP8.5 (high emission) scenarios. The shaded region shows the ensemble members' 10-90th percentile ranges.

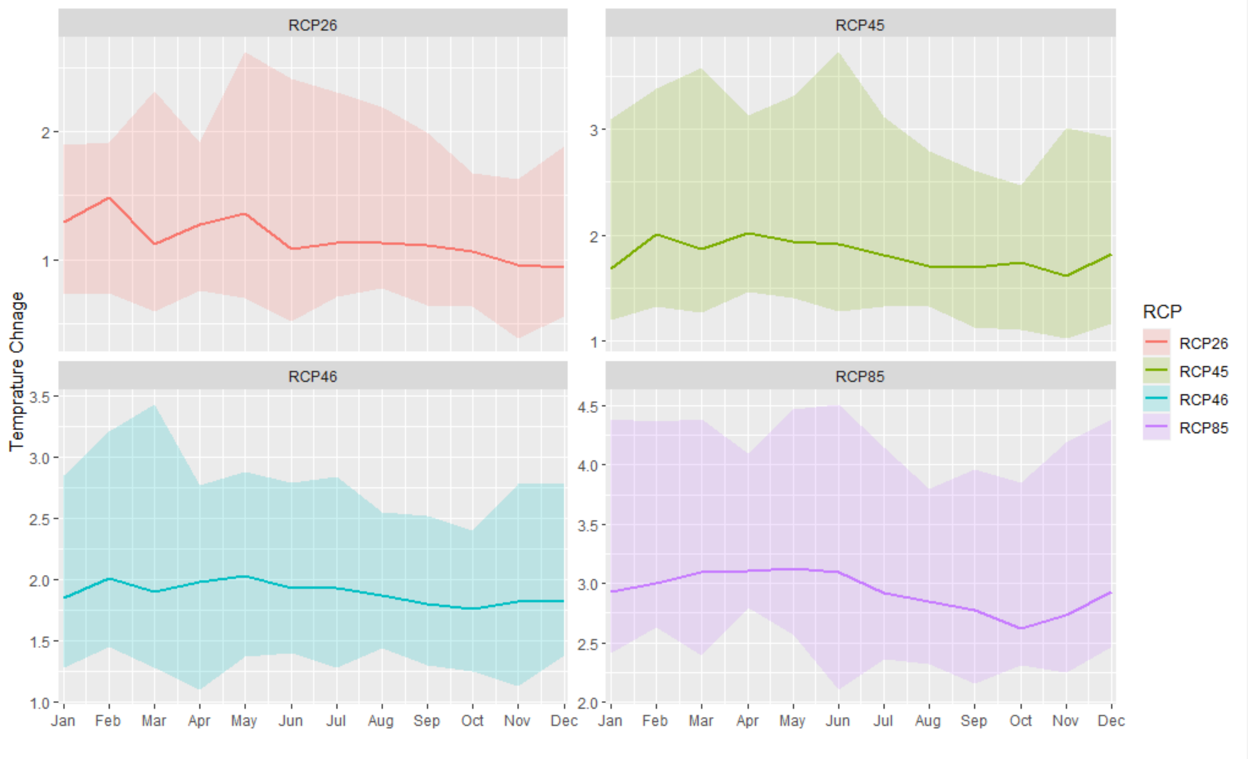


**Figure 4–figure supplement 7.** Mean change in monthly temperature compared to the reference period in the south Lake Tana (36.5-37.75 east, 10.7- 12 north) by 2070s under RCP2.6 (low emissions), RCP4.5 (medium low-emission), RCP6.0 (medium-high emission), and RCP8.5 (high emission) scenarios. The shaded region shows the ensemble members' 10-90th percentile ranges.

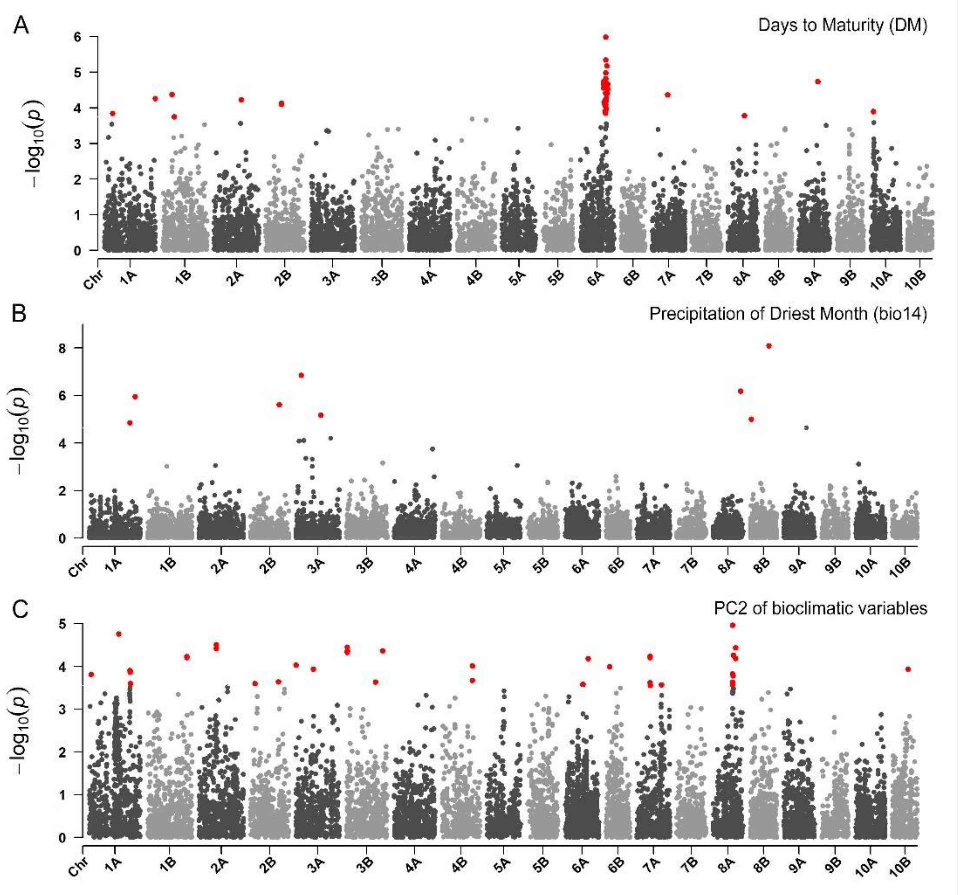


**Figure 5–figure supplement 1.** Manhattan plots reporting the GWAS result for (**A**) days to maturity, (**B**) precipitation of the driest month, and (**C**) PC2 of bioclimatic variables. On the x axis, the genomic position of markers. The y axis reports the strength of the association signal. SNPs are ordered by physical position and grouped by chromosome. Quantitative trait nucleotides (QTNs), *e.g.* SNPs surpassing a threshold based on a False Discovery Rate of 0.05, are highlighted in red.

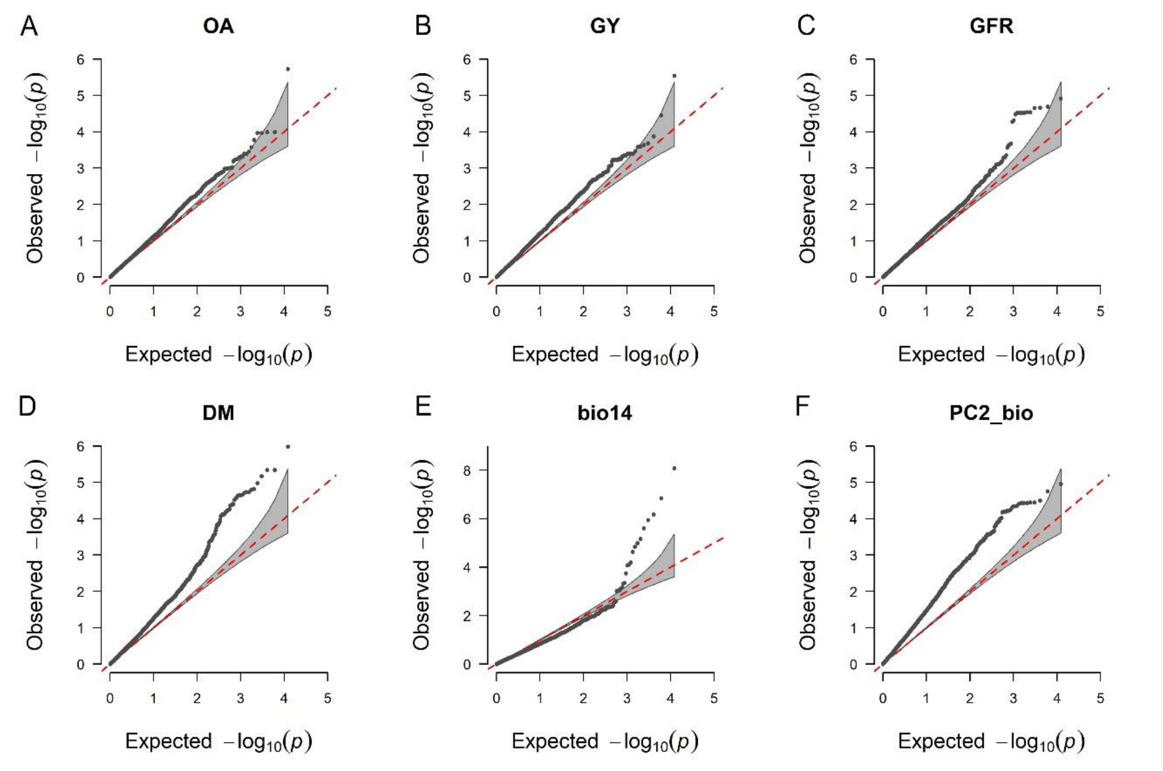


**Figure 5–figure supplement 2.** Quantile-quantile plots for the GWAS scans reported in Fig. 5 (A-C) and Fig. 5 supplement 1 (D-F). On the x axis, the expected distribution of p-values according to the null hypothesis of no association. On the y axis, the observed distribution of p-values. Each point represents an individual statistical test.
